# Deciphering the transcriptomic landscape of pyrazinamide resistance in *Mycobacterium tuberculosis*

**DOI:** 10.1101/2025.10.13.681998

**Authors:** Ananthi Rajendran, Ahmed Kabir Refaya, Kannan Palaniyandi

**Author notes:** **Address for Correspondence** Dr. Kannan Palaniyandi, Scientist “E,” Department of Immunology, ICMR-National Institute for Research in Tuberculosis, #1, Mayor Sathyamoorthy Road, Chetpet, Chennai 600031, India., **Email:**, **Institutional **.

## Abstract

Pyrazinamide resistance in *Mycobacterium tuberculosis* poses a major challenge to tuberculosis management and is primarily linked to *pncA* mutations, while the broader transcriptional adaptations underlying this resistance remain unclear. In this study, we performed a comparative analysis of the transcriptomic response of a mutant clinical strain and the drug-susceptible laboratory strain H37Rv under PZA exposure and non-exposure conditions. The clinical strain used for this analysis carried a 10-nucleotide deletion mutation in *pncA* (positions 118-127) that abolished PZA activation, identified in our previous study. The critical drug concentrations were established at 200 µg/mL for the clinical strain (CST) and 12.5 µg/mL for the H37Rv strain (RvT), with the untreated H37Rv strain (UTRv) used as a reference. RNA-sequence profiles from treated and untreated conditions were analyzed to identify differentially expressed genes, followed by functional enrichment, KEGG pathway mapping, and protein-protein interaction network analysis.

Analysis revealed 3,413 differentially expressed genes (padj ≤ 0.05), including 1,428 upregulated and 1,360 downregulated genes. Functional enrichment was predominantly detected in CST vs. RvT followed by CST vs. UTRv, whereas no significant enrichment emerged in RvT vs. UTRv, underscoring the mutation’s dominant influence on the pyrazinamide response. The ribosomal machinery genes *rplC, rplD, and rpsH* were significantly enriched and strongly upregulated in the mutant strain under treatment but only mildly regulated in the laboratory strain. We observed that several anti-TB drug targets (*katG, ethA, atpE, panD*) were downregulated, and a few efflux pumps (*Rv1258, Rv3008, Rv3756c*) were upregulated, reflecting cross-resistance mechanisms.

Network analysis identified 19 clusters, and prominent modules comprised polyketide synthases, PDIM synthesis genes, fatty acid β-oxidation enzymes and ESX secretion system. These interconnected modules highlight the metabolic and structural strategies that support persistence under drug pressure. Collectively, our findings link mutation-driven and PZA-induced transcriptomic alterations to adaptive pathways and provide insights into the mechanisms underlying tolerance and potential therapeutic opportunities.

**Author summary:** Pyrazinamide is a vital anti-tuberculosis drug, which helps to shorten the treatment duration and while resistance is common, it is not completely understood. Our previous work identified a clinical strain of *Mycobacterium tuberculosis* that was resistant to pyrazinamide due to a mutation in target gene responsible for drug activity. In this study, we compared the gene expression pattern of this resistant strain with a standard laboratory strain under drug exposure. The resistant strain exhibited distinct gene expression patterns, specifically, the activated genes associated with cell damage repair, lipid biosynthesis for the cell envelope, and ATP maintenance for energy production while it repressed genes that promote dormancy and virulence. Our findings suggest that the resistant strain is actively and metabolically adapts to the drug stress, rather than simply becoming dormant. We also found notable gene alterations in the other anti-tuberculosis drug targets, indicating possible cross resistance with pyrazinamide. Collectively, our findings provide insights into how genetic mutations alter the gene expression of resistant strains that adapt to survive drug exposure.

## 1. Introduction

Tuberculosis (TB) remains a global health concern in *Mycobacterium tuberculosis (M. tuberculosis)*, with an increasing burden of multidrug-resistant (MDR) and extensively drug-resistant (XDR) TB (1). Among the first-line TB regimens, pyrazinamide (PZA) is vital for eradicating semi-dormant bacilli and reducing the duration of TB therapy (2). However, the MDR-TB cases frequently exhibit PZA resistance (3), diminishing its clinical efficacy and requiring further research on the resistance mechanisms. The primary cause of resistance is attributed to *pncA* mutations, which encode pyrazinamidase (PZase) enzyme that converts PZA into pyrazinoic acid (POA) which is the active bactericidal form (4). However, evidence from our review emphasized that *pncA* mutations alone may not fully explain PZA resistance, and multiple alternative mechanisms have also been reported (5). Multiple reports have proven that broader transcriptional regulators, general stress responses, and metabolic pathways induce the multifaceted nature of drug resistance (Shi et al., 2020; Thiede et al., 2021).

Transcriptomic and other omics-based researches have been established to understand the Mycobacterial responses under drug pressure. Rifampicin-resistant strains have revealed significant alterations in gene expression patterns associated with *rpoB* mutations and compensatory regulatory mechanisms were reported (6). Capreomycin-resistant strains had loss or mutation in the *tlyA* gene which showed changes in the pathways involved in stress response and protein biosynthesis (7). When the MDR-TB strains treated with fluoroquinolone and ethionamide, the distinct transcriptional profiles of lipid metabolism, efflux mechanisms, and cofactor metabolism were reported. The altered expression of *Rv3094c* and downregulation of type VII secretion system components have also been observed in the ethionamide metabolism (8,9). Genome-wide transcriptional analysis has reported substantial modifications in transcriptional regulation and cell wall-related processes of streptomycin-resistant strains (Z. Wu et al., 2024). Similarly, the expression of PZA resistance-related genes, including *pncA*, has highlighted the regulatory framework, *rpsA* mutations that may result in reduced ribosomal binding to POA, potential efflux pumps, and *panD* mutations that could affect metabolic pathways (10,11).

A recent study reported that activation of the *sigE*-dependent cell envelope stress response plays a vital role in regulating PZA susceptibility, implying that sigma factor-mediated stress responses directly influence drug efficacy (12). *Mycobacterium tuberculosis* encodes 13 sigma factors, more than any other obligate pathogen per megabase of the genome, facilitating highly dynamic transcriptional responses to environmental and chemical stressors. Although *sigA* is essential for viability, alternative sigma factors are selectively activated under specific conditions, such as antibiotic exposure (13). Drug therapy induces stress responses in bacteria, and the degree of stress generated by antimicrobials has been shown to correlate with their efficacy (14). Consequently, sigma factors may play a key role in mediating both innate and acquired drug resistance by regulating the adaptive transcriptional programs that support survival under drug-induced stress (12).

Collectively, these studies have emphasized the drug resistance in *M. tuberculosis* is characterized by extensive transcriptional remodeling and stress induced adaptations. However, the specific transcriptomic effects of *pncA*-mediated PZA resistance remain poorly characterized. Understanding how these mutations alter gene expression under drug pressure is vital for identifying compensatory and adaptive pathways that support resistance. In this study, we have compared the transcriptome profile of this mutant clinical strain with susceptible reference strain, H37Rv. The PZA-resistant clinical strain used for this analysis harbouring a deletion of 10 nucleotides from position 118-127 in the *pncA* hindering the production of the *pncA* causing resistance was identified in our previous study (15). Post transcriptional analysis including differential gene expression (DEG) analysis, functional enrichment (FE), protein-protein interaction (PPI) network construction and pathway analysis was performed to elucidate the key genetic and regulatory mechanisms underlying PZA resistance and link these transcriptomic alterations to phenotypic drug tolerance.

## Results

### Characteristics of the strains and determination of critical drug concentration

The critical concentration of PZA for CST was 200 µg/ml and 12.5 µg/ml for RvT. The strains were grown in triplicates and RNA was extracted from each separately. As for the untreated control strain, RNA sequence from drug-free H37Rv strain was downloaded from (PRJNA579441) was termed as UTRv was used a drug free control for comparison (16). Details of all the strains used in the study are listed in Table 1. The BAM files of the PZA-treated resistant clinical strains CST1, CST2 and CST3 and PZA-treated H37Rv strains RvT1, RvT2 and RvT3 aligned with reference strain H37Rv was viewed in IGV and the presence of mutation at the locus (2289115 - 2289124) was confirmed in all the resistant strains, whereas no mutation was observed in reference strains.

**Table 1.** list of *M. tuberculosis* isolates.

| Strains | Condition |
| --- | --- |
| PZA treated *resistant* clinical strain _1 | CST1 |
| PZA treated *resistant* clinical strain _2 | CST2 |
| PZA treated *resistant* clinical strain _3 | CST3 |
| PZA treated H37Rv_1 | RvT1 |
| PZA treated H37Rv _2 | RvT2 |
| PZA treated H37Rv_3 | RvT3 |
| Untreated H37Rv_1 | UTRv1 (16) |
| Untreated H37Rv_2 | UTRv2 (16) |
| Untreated H37Rv_3 | UTRv3 (16) |

### Comprehensive transcriptome profiling across the three groups

Principal component analysis (PCA) of the normalized expression data revealed a clear separation of the three experimental groups along PC1 (75% variance) and PC2(23% variance) together capturing 98% variability in the dataset **(Fig 1A)**. Replicates from the same group clustered tightly, indicating high reproducibility, whereas distinct groups were clearly separated reflecting the major transcriptional differences induced by drug treatment. The sample-to-sample Euclidean distance heatmap further supported these distinctions, with shorter distances observed within biological replicates compared to between groups confirming high similarity in transcriptomic patterns within groups and marked divergence between groups **(Fig 1B)**.

**Fig 1.**
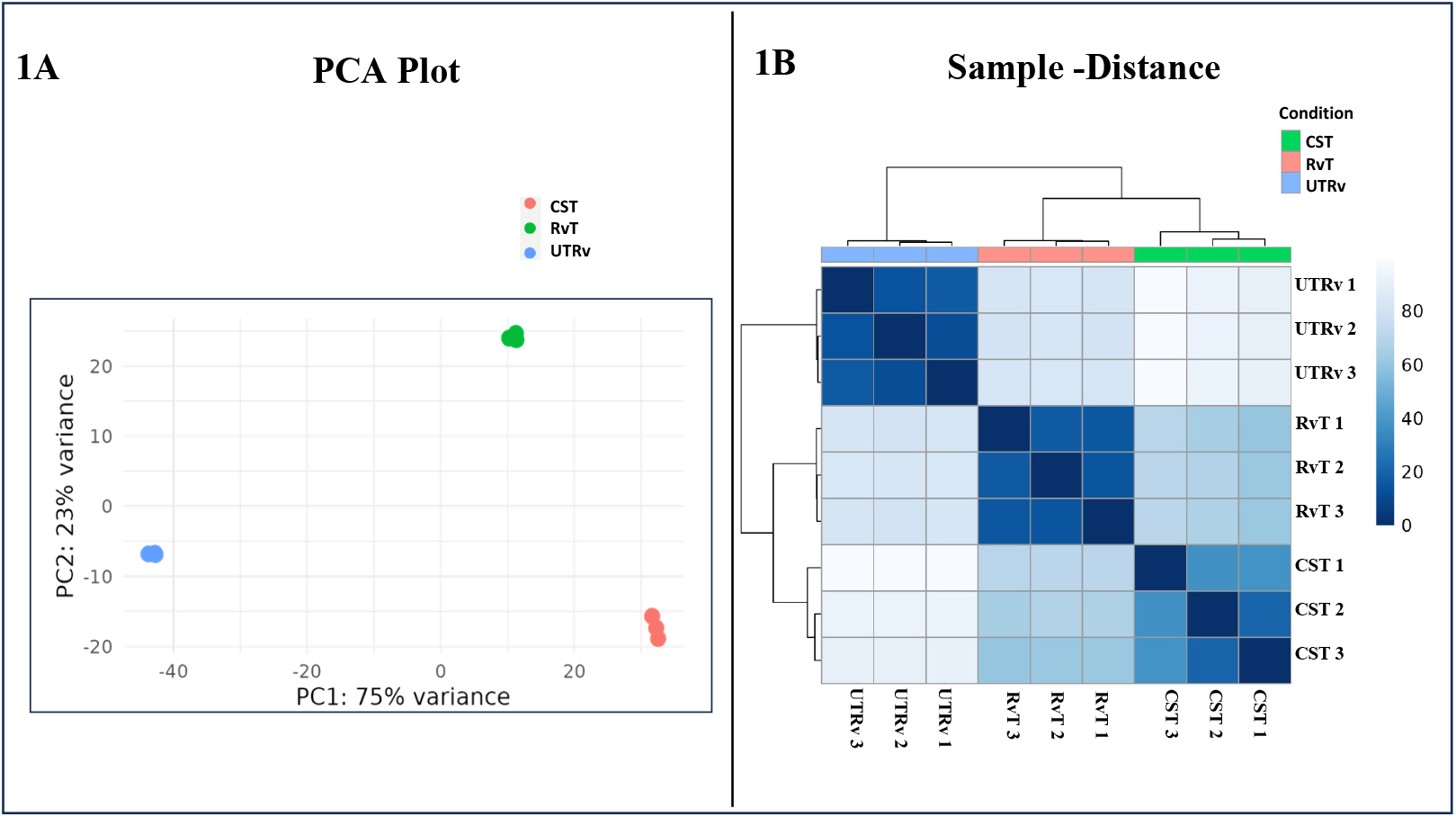
Transcriptomic variations among the experimental groups. (1A) Principal Component Analysis (PCA) of normalized expression data showed clear separation of the untreated control (blue), treated control (green), and treated mutant (red) groups. (1B) The pairwise distance between the genomes is depicted in the Euclidean heatmap. Darker shades indicate high similarity while the lighter shades indicate dissimilarity.

### Drug induced transcriptional changes in the laboratory strain

To investigate the transcriptional response of the RvT strain upon drug exposure, RNA-seq was used to compare its gene expression profiles with those of the UTRv. Differential expression analysis revealed a total of 2694 differentially expressed genes (DEGs) including 788 upregulated genes and 840 downregulated genes upon drug treatment, while 1067 genes were mildly regulated (-1< log2Foldchange < 1) but yet significant (padj≤0.05). The overall distribution of DEGs along with the top 10 most upregulated and downregulated genes is illustrated in the volcano plot **(Fig 2A)**, highlighting a significant increase in the expression of genes such as Rv3229 (DesA3) and Rv2947c (pks15), which are involved in lipid biosynthesis, and Rv2591(PE_PGRS44), a member of the PE/PPE family, which is implicated in host-pathogen interactions and immune modulation, while Rv2988c (leuC) encodes a probable 3-isopropylmalate dehydratase, which is important for amino acid biosynthesis. Several genes, such as Rv2625c, Rv2623, Rv1737c, Rv2032, Rv2628, and Rv2627, which are essential for *M. tuberculosis* survival under hypoxia, nitric oxide, and other conditions that induce dormancy, were significantly downregulated (17). Hierarchical clustering of the top 50 DEGs clearly distinguished treated from untreated samples **(Fig 3A)** highlighting a substantial transcriptional reprograming in response to drug exposure.

**Fig 2.**
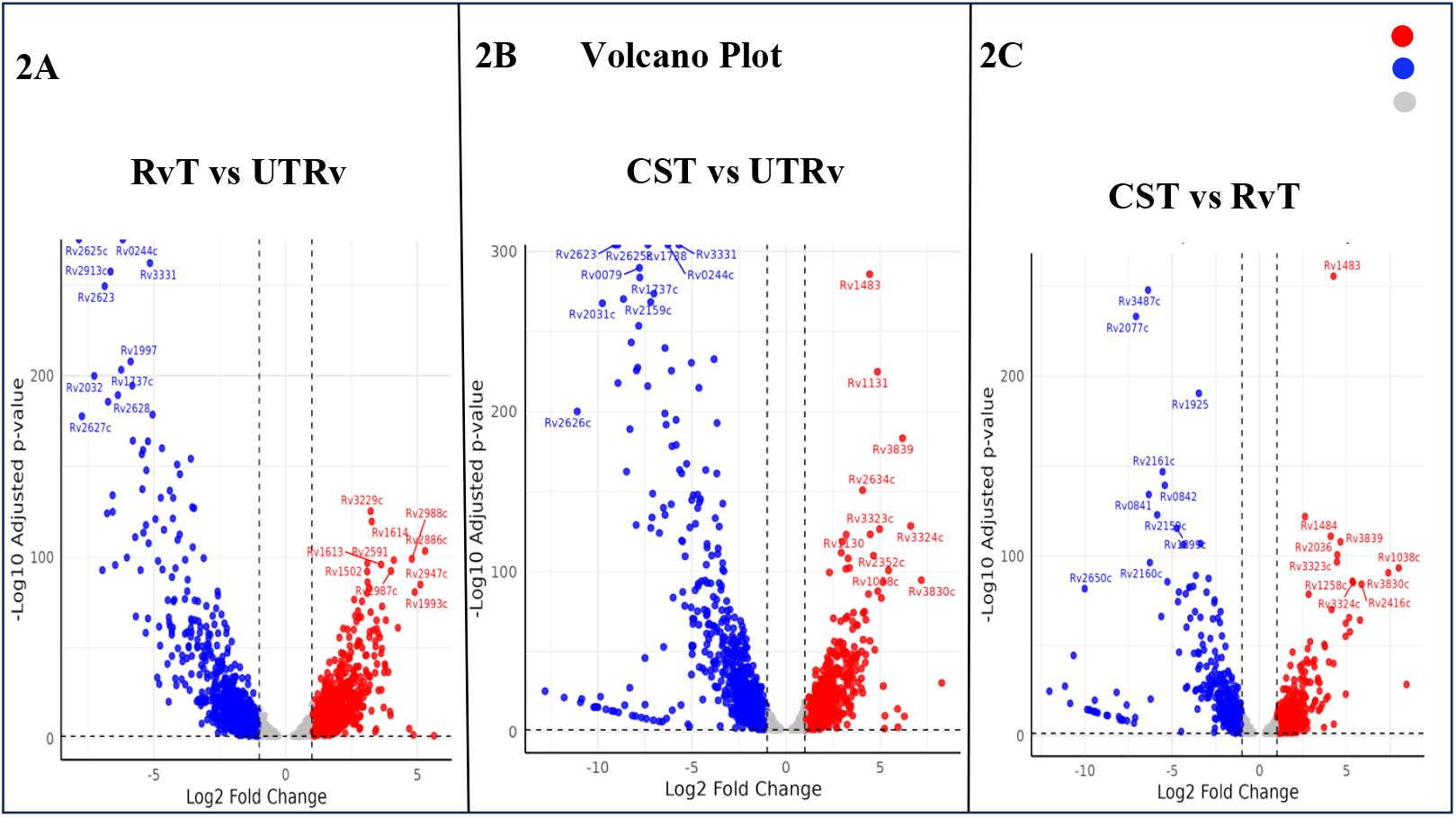
Volcano plots showing DEGs in three pairwise comparisons, 2A. RvT vs UTRv, 2B. CST vs UTRv and 2C. CST vs RvT. The x-axis shows log_2_ fold change and y-axis shows -log_10_ adjusted p-values. Horizontal and vertical dashed lines represent the thresholds for the adjusted p value (0.05) and log_2_ fold change (±1), respectively. Each point represents an individual gene, with red indicating significantly upregulated genes, blue indicating significantly downregulated genes, and grey indicating non-significant genes. The top DEGs based on statistical significance are labelled.

**Fig 3.**
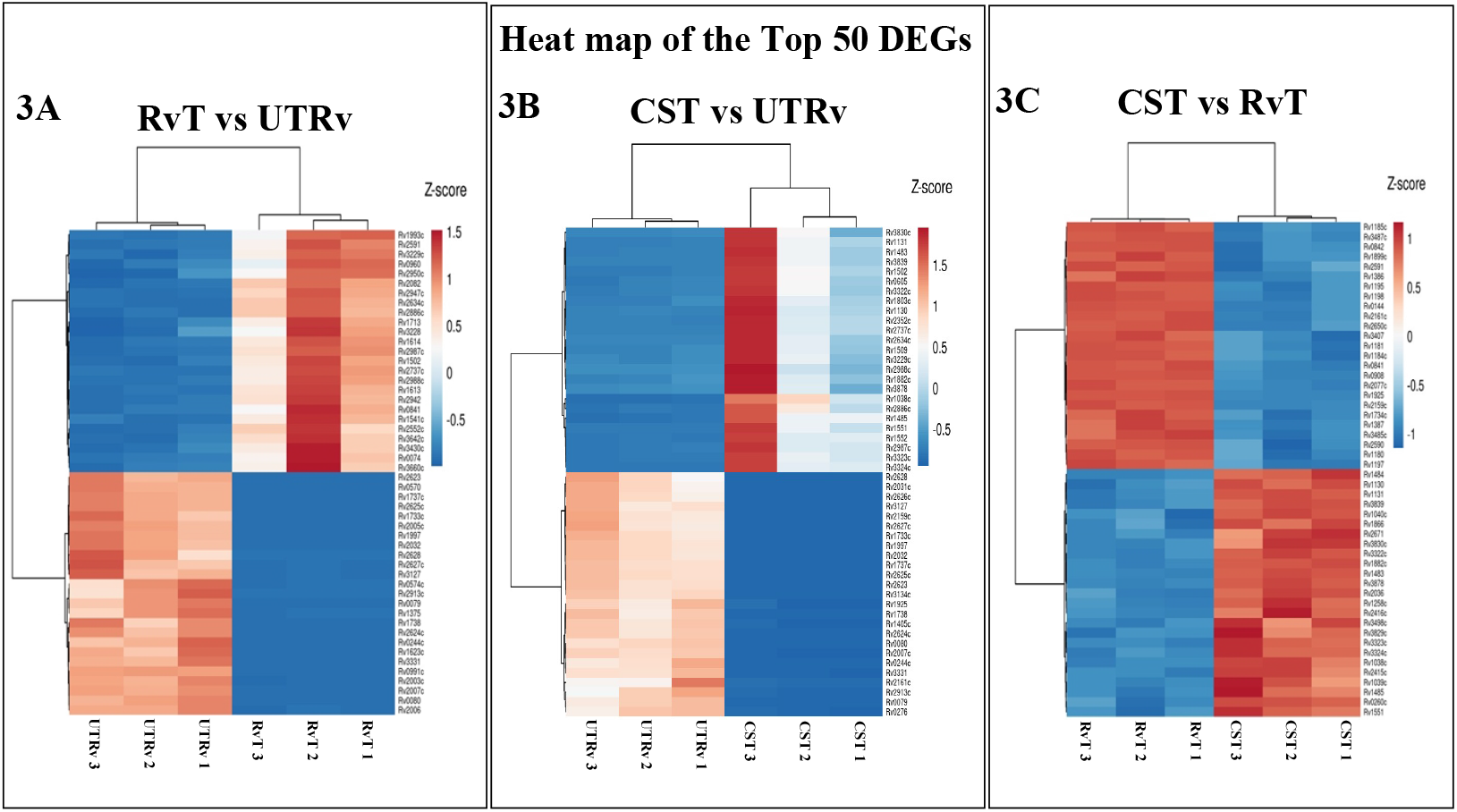
Heatmaps of top 50 differentially expressed genes: The heat map represents the expression pattern of top 50 differentially expressed genes across all samples. Gene expression values were normalized and scaled using z-scores. The rows represent the genes and the columns represent individual samples. The colour gradient indicates differential expression levels, with red indicating higher expression and blue indicating lower expression.

### Combined effect of mutation and drug treatment on mutant strain

In order to explore the dual effect of the mutation and drug pressure, the gene expression profiles of CST and UTRv were compared. About 2706 genes were differentially expressed including 832 upregulated genes and 827 downregulated genes and 1047 mildly regulated ones as depicted in the volcano plot **(Fig 2B)**. Among the top 10 genes, Rv1483 (fabG1), a β-ketoacyl-ACP reductase involved in mycolic acid biosynthesis; Rv1131 (prpC), involved in propionate detoxification; Rv3323c (moaX) and Rv3324c (moaC3), involved in molybdenum cofactor biosynthesis; and Rv3830c (TetR) transcriptional regulatory proteins were significantly upregulated. This suggests that the mutant strain responded to drug treatment by activating pathways related to cell wall maintenance, stress adaptation, and metabolic detoxification. Genes from the DosR regulon, such as Rv2623, Rv2626c, Rv2031c, and Rv1737c, which are known to be essential for dormancy and adaptation to hypoxia and nutrient starvation stress (17,18) and some conserved hypothetical genes such as Rv2625, Rv2159c, and Rv1738 were significantly downregulated. The heat map **(Fig 3B)** depicts the hierarchical clustering of the top 50 genes. The PZA treated mutant strain displayed a distinct transcriptional profile featuring suppression of stress adaptation pathways and activation of metabolic and repair-related genes suggests a compensatory shift that may mediate PZA resistance in *M. tuberculosis*.

### Distinct transcriptional responses of the clinical strain and laboratory strain to drug treatment

The gene expression profiles of CST and RvT were compared to identifying the effect of mutation on the gene expression. The comparison revealed a set of 566 upregulated genes, 487 downregulated genes and 1113 mildly regulated genes comprising a total of 2166 differentially expressed genes. The overall expression of the DEGs showing top 10 differentially expressed genes and the hierarchical clustering of the top 50 regulated genes are represented in the volcano plot **(Fig 2C)** and heatmap **(Fig 3C)** respectively. Notably, significant genes such as Rv1483 and Rv1484 (mycolic acid biosynthesis and cell wall maintenance), Rv2036 (stress adaptation), Rv3839 (conserved hypothetical protein), and Rv1038c (ESAT6 family genes) were highly expressed. This indicated that the CST strain may undergo enhanced cell envelope remodelling as a protective response, activation of additional stress adaptation, and metabolic pathways, potentially supporting bacterial persistence under PZA treatment. Conversely, genes including Rv3487c (lipid metabolism and host interaction), Rv2077c (cell wall/cell process protein), Rv1225 and Rv2161c (putative regulatory proteins), and Rv0842 (efflux pump linked to drug resistance) were downregulated. This suggests the suppression of certain virulence factors, cell wall processes, and drug efflux mechanisms, possibly as a trade-off to prioritize survival and adaptation under the selective pressure of PZA.

### Unique and Shared DEGs across the three groups

A total of 3413 (padj ≤ 0.05) DEGs were identified across three comparisons. However, when the overlaps were examined separately for strongly upregulated (log_2_FC ≥ 1 & padj ≤ 0.05) and downregulated (log_2_FC ≤ -1 & padj ≤ 0.05) genes, we observed 1428 DEGs were upregulated and 1360 DEGs were downregulated. CST vs. RvT exhibited 522 unique DEGs (294up, 228down) reflecting mutation associated transcriptional changes, whereas 254 unique DEGs (129up, 125down) were observed in CST vs. UTRv representing gene expression alterations primarily driven by drug treatment and 607 DEGs (302up, 305down) uniquely in RvT vs. UTRv indicating expression changes attributable to drug treatment on the control strain. A core set of 147 DEGs (55up, 92down) was found across all three comparisons, representing a shared transcriptional response to mutations and treatment conditions **(Fig 4A)**.

**Fig 4.**
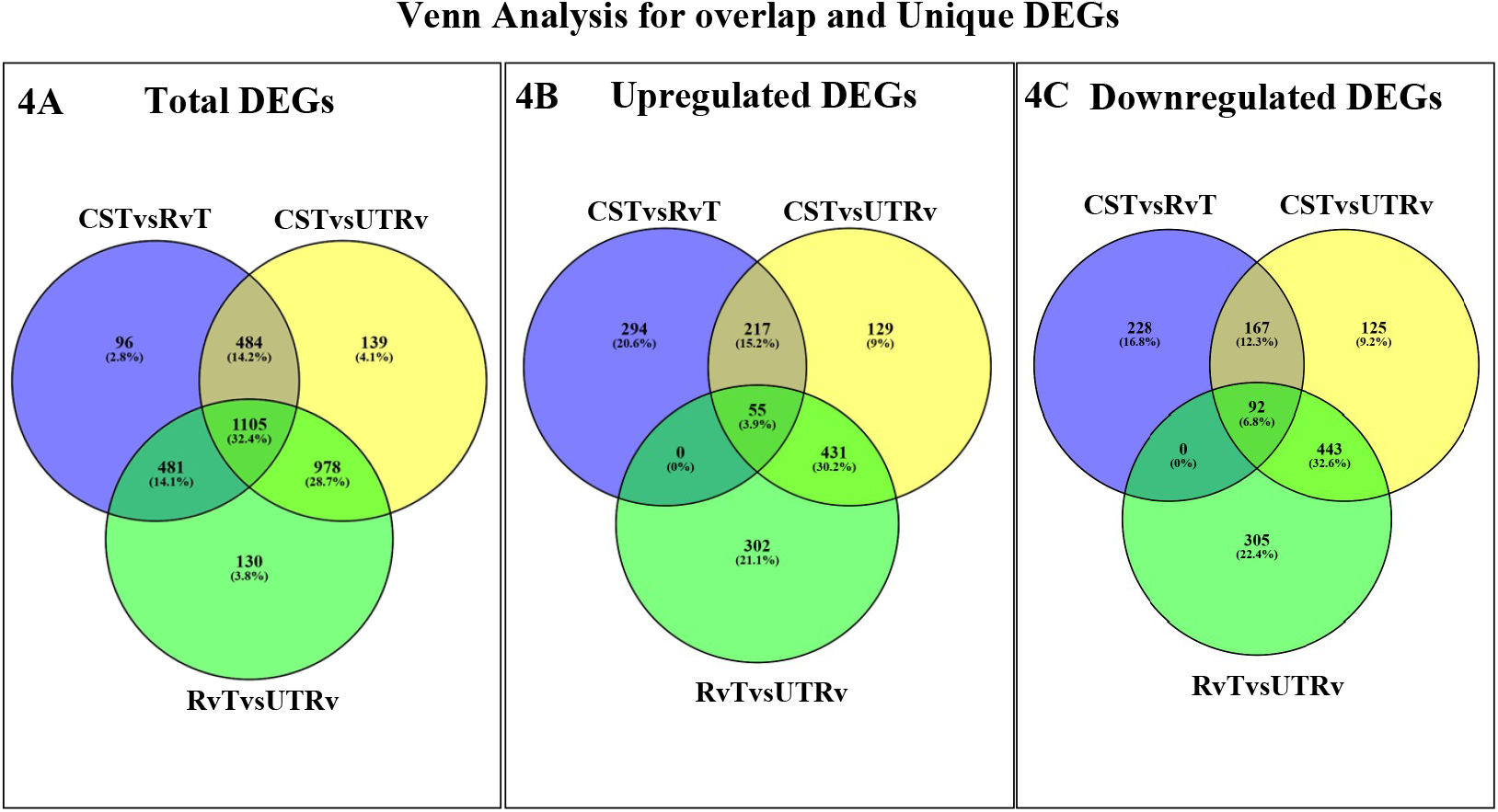
Venn diagram showing shared and unique genes across the three groups: The venn diagram illustrates the overlap of DEGs identified across each comparison. Each circle represents the set of DEGs from one comparison. Overlapping regions indicate genes shared between two comparisons, while non overlapping areas represent unique genes specific to each comparison.

### Venn Analysis for overlap and Unique DEGs

Between CSTvsRvT and CSTvsUTRv, 384 DEGs (217up, 167down) were shared likely reflecting mutation-specific gene expression signatures that persist even in the presence of drug pressure. CSTvsUTRv and RvTvsUTRv shared 874 DEGs (431up, 443 down) representing drug-responsive genes common to both the drug treated laboratory strain and mutant strain compared to the untreated control, reflecting conserved core pathways activated by PZA treatment. No genes met the criteria for significant up or down regulation between CSTvsRvT and RvTvsUTRv **(Fig 4B and 4C)**. This indicates that the 481 shared DEGs observed in the total DEGs were either upregulated or downregulated in either one of the comparisons rather in both or mildly regulated in at least one of the comparisons (-1 ≤ log_2_FC ≥1) despite being statistically significant (Padj < 0.05). This pattern reflects subtle but consistent transcriptional adjustments rather than strong directional shifts indicating variable regulation across conditions that reflect both common and specific responses.

### Biological features in functional enrichment and KEGG pathway

Functional enrichment and pathway analysis was performed separately for DEGs from each pairwise comparisons using DAVID. Both the analysis revealed a variable set of functionally enriched genes in CSTvsRvT, followed by CSTvsUTRv, while no enrichment was observed in RvTvsUTRv. Among all the enriched pathways and gene ontology identified, only one KEGG pathway and two molecular function terms reached statistical significance (FDR<0.05) especially in CSTvsRvT comparison. Specifically, the mtu03010: Ribosome pathway (FRD = 0.0065; 50 genes) and the molecular function term structural constituent of ribosome - GO:0003735 (FDR = 0.009; 50 genes), while the term rRNA binding – GO_0019843 showed borderline significance (FDR = 0.051; 31 genes). In addition to these, several other KEGG pathways and GO terms including GO:0006412; GO:0001666; GO:0006633; GO:0051607 were enriched at nominal p-values but did not reach the FDR<0.05 threshold. These have been listed in the **S1 Table** to provide a completer overview of all enrichment results. To visualize the distribution of the enriched pathways, a bubble plot was generated **(Fig 5)**, where each bubble represents an enriched pathway, bubble size reflects the number of genes involved and bubble colour indicates the FDR-adjusted significance level.

**Fig 5.**
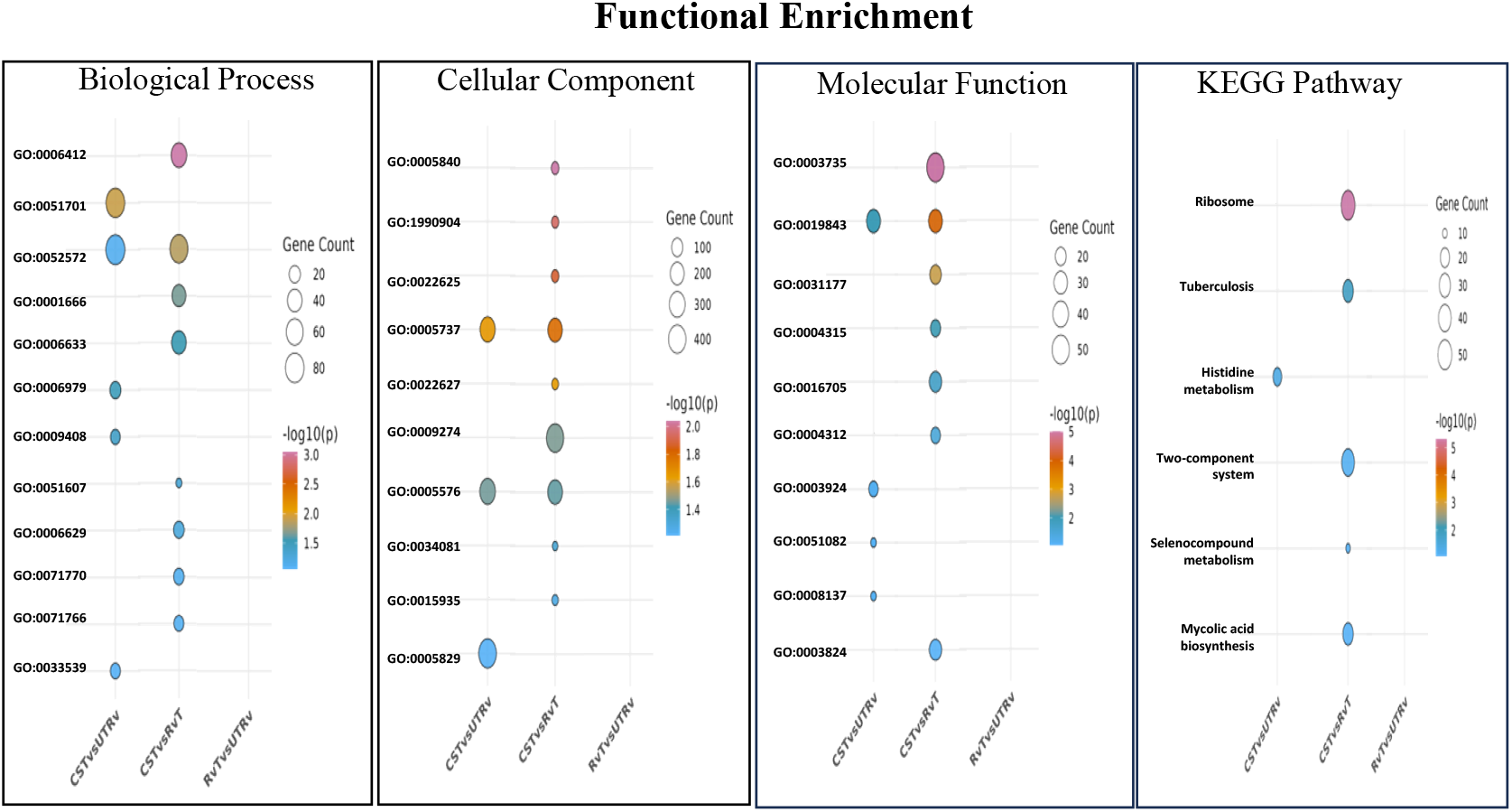
Functional enrichment analysis of DEGs in all three groups. The bubble plot depicts significantly enriched Gene Ontology (GO) terms including (A) Biological Process, (B) Cellular Component and (C) Molecular Function along with (D) KEGG pathways associated with DEGs. The colour gradient represents the statistical significance (–log_10_ *p*-value) of enrichment, while the bubble size corresponds to the number of genes associated with the respective GO term or pathway.

### Insights from the protein-protein interaction network

Among the 778 DEGs enriched in INTERPRO domains, 448 protein-coding genes were identified and used to construct a PPI network in STRING (v12.0) at high confidence threshold (> 0.7). The resulting network comprised 445 nodes and 1286 edges, with an average node degree of 5.78 and an average local clustering coefficient of 0.44. The observed PPI network showed significant enrichment (p-value < 1.0e-16), indicating that the proteins are functionally connected rather than randomly associated. MCODE clustering of the PPI network identified 19 distinct clusters, representing densely interconnected protein modules. Cluster 1 was the largest containing 25 genes followed by cluster 2 with 15 genes, cluster 4 with 14 genes, cluster 12 with 12 genes, cluster 3 with 10 genes and cluster 6 with 6 genes. The remaining clusters contained five or fewer genes. Functional annotations of these clusters revealed enrichment of multiple biological processes, cellular component, molecular functions and KEGG pathways. Some genes were assigned to a single GO term while others were annotated to several terms resulting in overlaps among functional categories (**S2 Table)**. To visualize the expression trends and interaction confidence, nodes were coloured according to their log2fold change values and the edge thickness was scaled according to the combined STRING score **(Fig 6)**.

**Fig 6.**
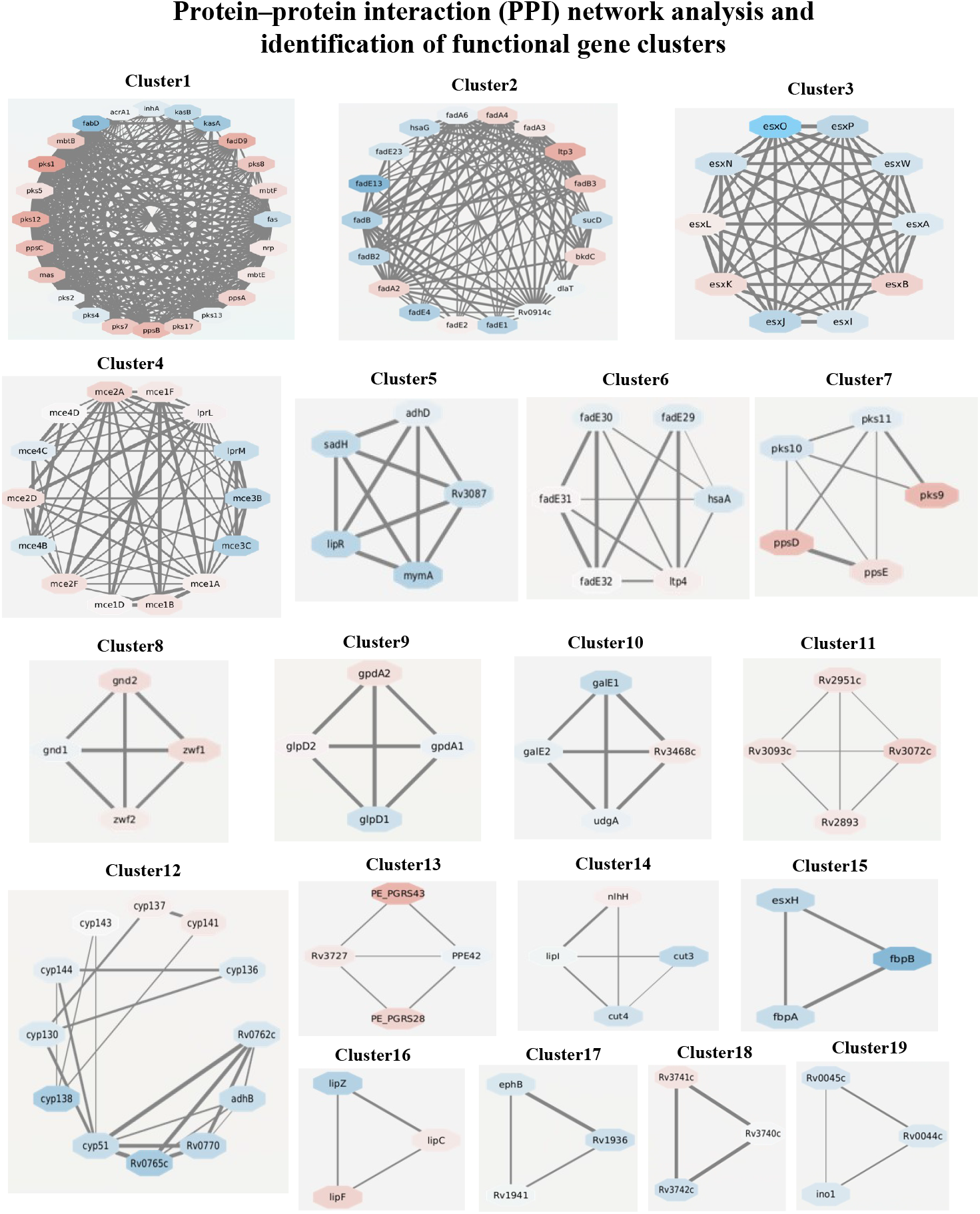
Protein–protein interaction network and functional gene clusters. A high-confidence PPI network (STRING v12.0; score > 0.7) was generated using protein-coding DEGs enriched in INTERPRO domains and yield 19 densely connected gene modules. Clusters 1–4 were the largest clusters consisting of gene modules associated with lipid metabolism, ESX secretion systems, and mce transport operons. Clusters 5-19 were smallest gene modules, but were still grouped in their functional coherence. The nodes are color coded by log_2_ fold change (red - upregulated; blue - downregulated) and the edge thickness corresponds to confidence in interactions, representing transcriptional variations and the connectivity in the network.

## DISCUSSION

This study provides comprehensive transcriptomic alterations associated with *pncA*-mediated PZA resistance in *M. tuberculosis*. We differentiated the transcriptomic profiles of a clinical strain harboring a 10-nucleotide deletion in *pncA* and the drug-susceptible laboratory strain H37Rv and identified distinct transcriptional signatures that elucidated how the resistant strain adapts to PZA stress. To our knowledge, this is the first study to report the whole transcriptome profile of a PZA-resistant clinical isolate of *M. tuberculosis* using a high-throughput RNA-sequencing genome-wide approach. This revealed the full transcriptional landscape and integrated changes in the metabolic, stress response, and ribosomal pathways. Thus, this high-resolution dataset offers a combined view of *M. tuberculosis* reprogramming and gene expression under the dual effect of *pncA* mutation and PZA exposure. The resistant mutant demonstrated a comprehensive alteration in gene expression, with marked upregulation of metabolic and cell envelope biosynthetic pathways and repression of genes associated with dormancy and virulence. These changes suggest that *M. tuberculosis* adopts a metabolically active state rather than a drug-induced dormant state to survive under drug pressure.

The PZA susceptible RvT strain produced only modest transcriptional shifts, with few genes meeting the significance threshold and no functional categories passing the enrichment cut-offs (RvT vs. UTRv). This indicates that short-term drug treatment of the laboratory strain mainly elicits a general stress response without activating specific adaptive pathways. In contrast, the CST strain exhibited a pronounced and distinct response compared to both UTRv and RvT strains. Notably, the upregulated genes of CST strain involved in lipid metabolism, mycolic acid biosynthesis and molybdenum cofactor synthesis, indicating that the mutant strain activates pathways linked to cell envelope maintenance, metabolic adaptation, and detoxification when challenged with PZA. Simultaneously, multiple members of the DosR regulon, certain virulence factors, phospholipases, efflux pumps, and other dormancy-associated genes were consistently downregulated. Such selective suppression of non-essential virulence functions and dormancy-associated genes suggests a compensatory reprogramming of the CST strain away from a latent-like state toward active metabolic repair, which may help sustain growth under drug pressure. These findings are consistent with previous observations in multidrug-resistant and extensively drug-resistant *M. tuberculosis* strains, which show altered lipid metabolism, cell wall dynamics, and stress responses under antimicrobial pressure (17–19).

Functional enrichment analysis demonstrated a clear enrichment pattern in the CST vs. RvT, followed by weaker but related signals in the CST vs. UTRv. In contrast, no pathways or GO terms reached significance in the RvT vs. UTRv comparison, indicating that PZA treatment of the laboratory strain did not elicit a coordinated functional response. Among the enriched terms in CST vs. RvT, the KEGG ribosome pathway (mtu03010) and the GO molecular function structural constituent of the ribosome (GO:0003735) were the most significant, with rRNA binding (GO:0019843) narrowly missing this threshold. This pattern suggests that under drug pressure, the resistant mutant alters the ribosomal and translational machinery components, possibly reflecting enhanced protein synthesis or compensatory remodelling of translation as part of its adaptive response. Almost 50 genes associated with the ribosomal machinery were found to be strongly enriched across all three categories in the mtu03010, GO:0003735, and GO:0019843, which represent the core ribosomal protein set. Most of these genes were common across all three ribosome-related terms, although a few additional rRNA-binding genes, Rv2364, Rv2925, Rv3462, Rv1165, and Rv2477, were uniquely identified under the rRNA-binding function. Prominent genes such as Rv0682, Rv0700, Rv0701, Rv0702, Rv0704, Rv0708, Rv0715, Rv0718, Rv1037, and Rv1038 were strongly upregulated, and Rv1197 and Rv1198 were downregulated in CST compared to both RvT and UTRv. Some genes, such as Rv0056 and Rv2347, were upregulated, and Rv1642 and Rv3874 were downregulated in CST vs. RvT but not in CST vs. UTRv. The difference in the expression levels of these genes between the two comparisons may reflect the different pressures involved. For instance, in CST vs. RvT, the changes observed are mainly driven by the mutation under drug treatment, whereas in CST vs. UTRv, the effects of both mutation and drug pressure are combined, leading to less pronounced regulation.

Studies reported that ribosomal machinery components are targets for many anti-tuberculosis drugs (20–22). The ribosomal subunits within PPI networks present promising targets for the development of novel antibiotics, owing to the structural differences between bacterial and human ribosomes (23). Moreover, the genes Rv0702 (rplD) and Rv0701 (rplC) are jointly inhibited by oxazolidinone-class antibiotics, including linezolid and sutezolid, highlighting the relevance of ribosomal components as drug targets (22). *M. tuberculosis* typically downregulates ribosomal protein genes during dormancy and under stress conditions, such as nutrient starvation, hypoxia, and antibiotic exposure, reflecting an energy-conserving survival strategy (24). Together, these findings indicate that the mutant strain favors an active metabolic survival strategy rather than a dormancy-driven strategy.

Furthermore, we observed differential expression of genes associated with other anti-TB drugs and their corresponding drug targets in CST vs. UTRv, suggesting possible implications for cross-resistance mechanisms. Notably, the drug targets for PZA (*pncA, panD*), isoniazid (*katG, ahpC*), ethionamide (*ethA*), delamanid (*fgd1, Ddn*), and bedaquiline (*atpE, Rv0678, mmpL5, mmpS5*) were also downregulated, suggesting a potential link between PZA resistance and resistance to other anti-TB drugs. This finding is consistent with the results of a previous study, which reported a high correlation between PZA resistance and resistance to streptomycin, ethambutol, ofloxacin, and levofloxacin, suggesting that the accumulation of genetic mutations in MDR-TB strains may contribute to cross-resistance (25). These studies clearly show that bacteria that develop resistance to one drug may modulate the expression of other drug targets to enhance their survival. Therefore, we hypothesized that the observed downregulation of these drug targets might be part of a broader cross-resistance mechanism. Additionally, we observed that the previously reported PZA resistance efflux pump genes, such as Rv3756c, Rv3008, Rv1258, and Rv0191 (26) and MDR-TB efflux genes *Rv2936, Rv2937*, and *Rv1643* (27) were upregulated, clearly indicating that the upregulated efflux pumps, which are proteins that actively expel a variety of antibiotics from the cell, thereby reducing the effectiveness of the drugs. These pumps can lead to cross-resistance among different antibiotic classes. The upregulation of efflux pumps and antibiotic-degrading enzymes in both CST and RvT aligns with the findings of studies on ethionamide, streptomycin, and fluoroquinolone resistance, in which broader stress and metabolic pathways have been implicated (8,9,28).

PPI network analysis revealed a complex interaction landscape comprising 19 distinct clusters, reflecting the multifaceted metabolic and virulence programs of *M. tuberculosis*. All clusters of genes identified in the PPI network and its functional enrichments were listed in **S2 Table**. The majority of the genes were predominantly grouped into five clusters. Cluster 1 integrated polyketide synthase genes, β-ketoacyl synthases, PDIM synthesis genes, and siderophore biosynthesis genes, highlighting the convergence of lipid biosynthesis and iron acquisition pathways that underpin mycobacterial adaptation and persistence (29,30). Cluster 2 was enriched with *fadA* and *fadE* family genes, reflecting the reliance on fatty acid degradation and β-oxidation for energy generation from host-derived lipids (31). Cluster 3 was dominated by ESX family genes, representing the Type VII secretion system (T7SS), which secretes key virulence factors critical for immune modulation and host-pathogen interactions (32,33). Cluster 4 represents a network of mce genes, underscoring the role of membrane transporters in lipid metabolism and host interactions, and identifying them as potential therapeutic targets (34). Cluster 12 was dominated by cytochrome P450 (CYP) genes, reflecting a specialized oxidative module involved in diverse redox reactions that are crucial for *M. tuberculosis* metabolism and adaptation (35). Taken together, the emergence of these distinct yet interconnected modules spanning lipid and iron acquisition, fatty acid degradation, pentose phosphate pathway activity, glycerol metabolism, polysaccharide biosynthesis, and CYP-mediated oxidative processes underscore how *M. tuberculosis* orchestrates metabolic and structural pathways to maintain its persistence under acidic, drug-challenged conditions. The emergence of these interconnected modules highlights how *M. tuberculosis* coordinates lipid metabolism, redox homeostasis, and virulence pathways, suggesting that targeting such functional hubs may offer novel therapeutic opportunities.

## Methods

### Growth and drug treatment of *M. tuberculosis* strains

Monoclonal cultures of the *pncA* clinical mutant strain and the laboratory strain H37Rv were used as the parental strain (P0) and were initially grown on drug-free 7H11 agar with fetal bovine serum (FBS) instead of oleic acid-albumin-dextrose-catalase (OADC) to support the substantial growth of *M. tuberculosis* at a lower pH. The medium was maintained at an acidic pH to retain the PZA activity which differs from the conventional pH of 6.8 (Heifets &Sanchez, 2000). The cultures were incubated at 37^0^C for 4 weeks to allow full growth. Following this initial state, the cultures were transferred to 7H11 agar containing PZA (Sigma-Aldrich,USA) at varying concentrations of 6.25 ug/ml, 12.5 ug/ml, 25 ug/ml, 50 ug/ml, and 100 ug/ml for the laboratory strain H37Rv and 100 and 200 ug/ml for the *pncA* clinical mutant strain. After four weeks of incubation, the critical concentration was determined for each strain and the strains that exhibited growth at or above this critical concentration were designated as P1 strains. These strains were further sub cultured onto fresh agar plate containing drug and incubated under similar conditions to generate P2 strains. The process of subculturing was repeated sequentially through P3 and P4 stages with each passage lasting four weeks at 37^0^C. Finally, the P4 strains which maintained growth at their respective critical concentration were selected for RNA extraction. The PZA-treated resistant clinical strain was designated as CST, the PZA-treated H37Rv strain as RvT.

### Transcriptome sequencing and analysis

Total RNA was extracted from each strain at three biological replicates with established protocols (36). RNA purity and quantity was estimated after the DNase I treatment using an Agilent 2100 Bioanalyzer. cDNA libraries were generated using Roche’s KAPA RNA Hyper Prep Kit as per the manufacturer’s instruction. The library quality was checked with an Agilent Bioanalyzer 2100 system and sequencing was performed in Illumina NovaSeq 6000 system. The raw read RNA-seq quality was assessed using FastQC, followed by low-quality bases and adapter sequences were removed using trimming and filtering with FastP. The processed reads were aligned to the reference genome (NC_000962) using HISAT2 v2.2.6 (37). The resulting BAM file was visualized using the Interactive Genomics Viewer (IGV) to check strain-specific mutations at the transcriptomic level.

### Comparative transcriptome Analysis

We have performed comparative transcriptomic analysis using RVT and CST strains along with untreated control strain obtained from previous study (16). The preliminary analysis and alignment were uniformly performed for all the datasets. All the datasets were processed identically using FastQC for quality control, FastP for trimming and HISAT2 for alignment to the reference genome. Gene-level read counts was achieved using feature counts (v 2.0.3) from the Subread package (38) and the Differentially Expressed Genes (DEGs) were identified using DESeq2 (v1.40.1) in R (39). Principal component analysis (PCA) and sample-to-sample distance plots were employed for data exploration and quality assessment. Genes with an adjusted p-value (FDR) <0.05 and log2fold change >1 and < -1 were considered significantly differentially expressed. Expression levels were quantified as Fragments per kilobase of transcript per million mapped reads (FPKM). Visualization of DEGs was performed in R using the enhanced volcano package for volcano plots (https://bioconductor.org/packages/release/bioc/html/EnhancedVolcano) and pheatmap package for clustered heat maps (https://cran.r-project.org/web/packages/pheatmap).

### Functional enrichment analysis

Differentially expressed genes from each pairwise comparison were used as input gene list for functional enrichment. All genes detected in the RNA-seq dataset were used as the background set. Enrichment analyses were carried out using the DAVID (Database for Annotation, visualisation and integrated discovery) Bioinformatics Resources 2021, (https://davidbioinformatics.nih.gov/tools.jsp) with *M. tuberculosis* H37Rv as the reference (40,41). The annotation categories, including Biological Process (BP), Molecular Function (MF), Cellular Component (CC) and KEGG pathways (KP) were analysed. Significance was assessed using the modified Fisher’s exact test and Benjamini-Hochberg FDR correction. Categories with FDR<0.05 were considered significantly enriched.

### Protein-protein interaction (PPI) network interaction

All the DEGs enriched in the INTERPRO domain of DAVID were used inputs for the STRING database (version 12.0, http://string-db.org) (42). Interactions were retrieved at a high confidence score of 0.7 and medium FDR stringency. Networks were imported into Cycoscape (version 3.10.2) for visualization and analysis (43). Clusters were identified using the MCODE plugin in Cytoscape with the default parameters. The resulting subnetworks were examined for their functional enrichment. Nodes were coloured according to the log2foldchange values and the edge thickness reflected the combined score.

## Supporting information

**S1 Table**. Functionally enriched DEGs from multiple annotation categories across three groups. This table summarizes the enrichment results of DEGs for different annotation categories includes COG ontology, Biological Process, Cellular Component, Interpro, Molecular Function, and KEGG pathway comprised in separate sheets. For each category, the columns include the comparison group, enrichment category, term identifier or description, number of genes in the term, p-value, list of genes contributing to the enrichment, fold enrichment, and multiple testing correction values (Bonferroni, Benjamini, and FDR). These results provide an overview of the key functional terms and pathways significantly associated with transcriptional changes under different experimental conditions.

**S2 Table**. Gene clusters derived from the protein–protein interaction network and their functional enrichments. This table contains three sheets representing Cluster 1, Cluster 2, and Other Clusters obtained from PPI network analysis. Each sheet lists the genes associated with the respective clusters along with their functional annotations. The table detailed about the clusters, node name, gene ID, protein ID, comparison group, functional category, and the GO or pathway terms that were enriched. This dataset highlights the functional organization of differentially expressed genes within key interaction modules.

**S1 Data**. RNA sequencing data associated with this study have been deposited in the NCBI BioProject under accession number PRJNA1151825. (https://dataview.ncbi.nlm.nih.gov/object/PRJNA1151825?reviewer=6vdhfan846tjpia4qfc3hls3ph)

## Acknowledgments

The authors would like to thank the Director of ICMR-National Institute for Research in Tuberculosis, for facilitating the execution of the study. Furthermore, the invaluable experimental assistance provided by Dr. S. Balaji and valuable inputs on data analysis by Dr. V. Umashankar is duly acknowledged.

## Author contributions

**Conceptualization:** Kannan Palaniyandi, Ananthi Rajendran.

**Data curation:** Ahmed Kabir Refaya, Ananthi Rajendran.

**Formal analysis:** Ananthi Rajendran, Ahmed Kabir Refaya.

**Funding acquisition:** Kannan Palaniyandi.

**Investigation:** Ananthi Rajendran, Kannan Palaniyandi.

**Methodology:** Ananthi Rajendran.

**Software:** Ahmed Kabir Refaya.

**Project administration:** Kannan Palaniyandi.

**Resources:** Kannan Palaniyandi.

**Supervision:** Kannan Palaniyandi.

**Validation:** Ahmed Kabir Refaya, Ananthi Rajendran.

**Visualization:** Ananthi Rajendran, Ahmed Kabir Refaya, Kannan Palaniyandi

**Writing – original draft:** Ananthi Rajendran.

**Writing – review & editing:** Ananthi Rajendran, Ahmed Kabir Refaya, Kannan Palaniyandi.

